# Critical residues of the antibiotic peptide Lys^M^ that inhibits lipid II flipping

**DOI:** 10.1101/2023.05.22.541694

**Authors:** Hidetaka Kohga, Napathip Lertpreedakorn, Ryoji Miyazaki, Tomoya Tsukazaki

**Affiliations:** Nara Institute of Science and Technology, Ikoma, Nara 630-0192, Japan

**Author notes:** Correspondence authors. (H.K.); (T.T.). These authors contributed equally (Hidetaka Kohga, Napathip Lertpreedakorn).

## Abstract

Small single-strand DNA/RNA phages that infect gram-negative bacteria encode lysis proteins that induce cell lysis without directly degrading the cell wall. One such protein, the 37-residue Lys^M^ protein derived from a lysis gene of *Levivirus* phage M (*lys*^*M*^), completely blocks the lipid II transport activity mediated by *Escherichia coli* MurJ, which is essential for peptidoglycan biosynthesis. Lys^M^ was proposed to be a single α-helical transmembrane protein that binds to MurJ and prevents its conformational transition during lipid II transport. Although Lys^M^ possibly interacts with MurJ, the inhibition mechanism remains unclear. Here, we identified the crucial residues for Lys^M^ function via comprehensive alanine-scanning mutagenesis. These residues were located on two surfaces in an α-helix model, probably providing surfaces interacting with MurJ in the membrane. This study provides fundamental information regarding the mechanism of Lys^M^ inhibition.

## Introduction

The emergence of multidrug-resistant bacteria is a major problem in public health. Thus, new types of antibiotics are urgently needed. In gram-negative bacteria, the peptidoglycan (PG) layer between the inner and outer membranes is a fundamental component with essential roles, such as maintaining cell shape and adapting to environmental changes (Silhavy, Kahne, & Walker, 2010; Sun, Rutherford, Silhavy, & Huang, 2022). Therefore, antibiotics are often designed to inhibit cell wall biogenesis.

The PG layer mainly comprises glycan chains with polymerized units consisting of N-acetylglucosamine (GlcNAc) and N-acetylmuramic acid (MurNAc). Tetrapeptides bind to MurNAc crosslinked to adjacent glycan chains, forming polymeric mesh structures in the periplasm (Figure 1A). The initial pathway of PG synthesis occurs on the cytoplasmic side. First, muramyl ligases (MurA-F) synthesize UDP-MurNAc-pentapeptide. Then MraY connects UDP-MurNAc-pentapeptide to undecaprenyl phosphate, generating lipid I (Und-PP-MurNAc-pentapeptide). Afterward, lipid I is converted into lipid II by MurG using UDP-GlcNAc. Lipid II, facing the cytoplasm, is flipped across the cytoplasmic membrane to the periplasm by the flippase MurJ (Inoue et al., 2008; Ruiz, 2008, 2015; Sham et al., 2014). Finally, lipid II is assembled into the PG layer after multiple steps, including polymerization.

**Figure 1.**
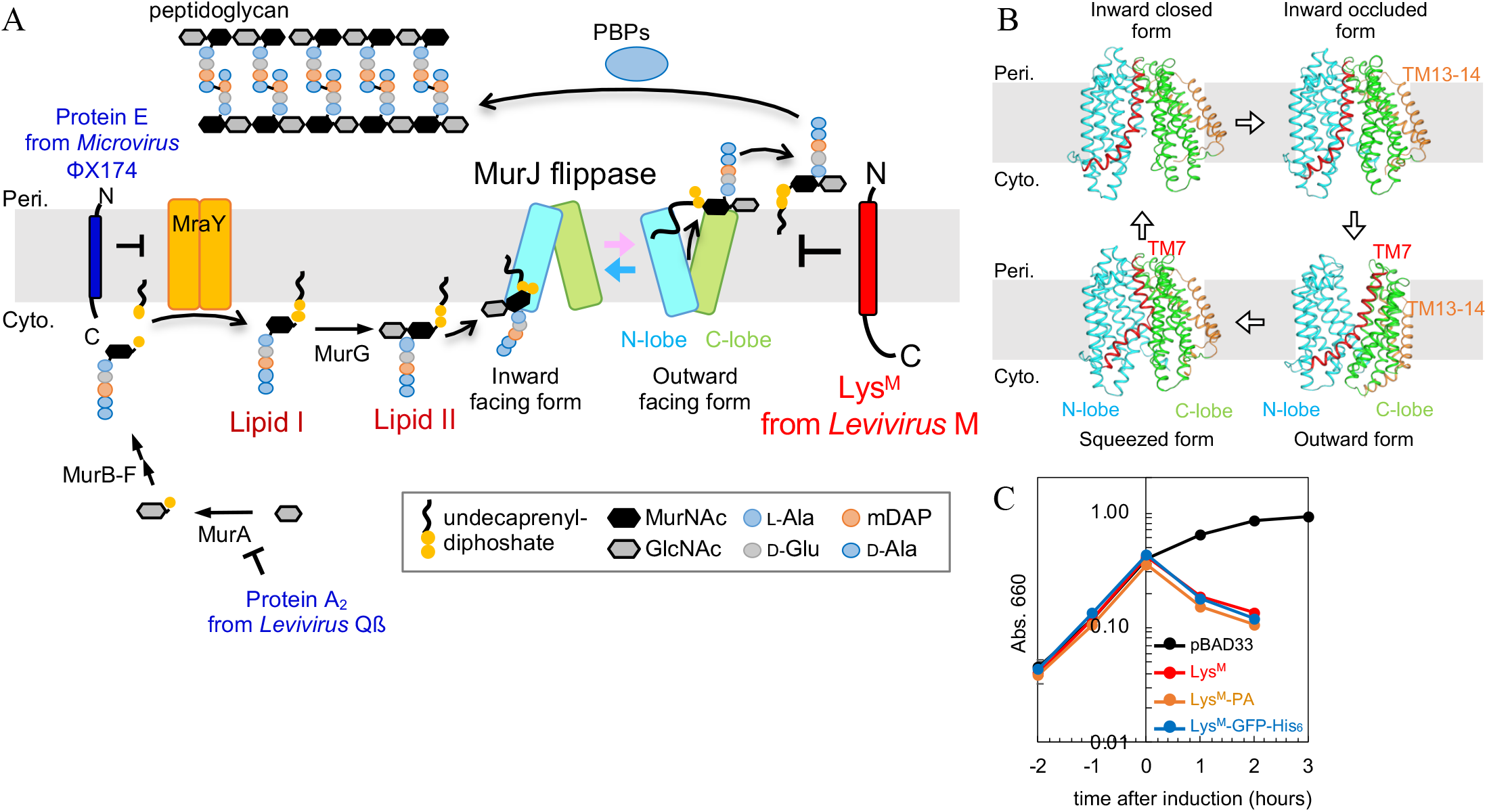
Inhibition model and *Escherichia coli* lysis by Lys^M^. (A) Peptidoglycan biosynthesis pathway. Lys^M^ is proposed to inhibit the flipping of lipid II by MurJ. (B) Previously proposed model of MurJ conformational changes. Cartoon models viewed from the membrane plane were generated using available MurJ crystal structures. PDB IDs of MurJ crystal structures: inward closed form of *E. coli* MurJ (PDB: 6CC4), inward-occluded form of *Arsenophonus endosymbiont* MurJ (PDB: 7WAX), outward form of *Thermosipho africanus* MurJ (PDB: 6NC9), and squeezed form of *E. coli* MurJ (PDB: 7WAG). The N-lobe, C-lobe, and additional helicase regions are colored blue, green, and orange, respectively. (C) *E. coli* cell lysis by Lys^M^. The growth of *E. coli* cells harboring pBAD33 (vector) or plasmids carrying the cloned *lys*^*M*^ genes is shown. Cells were grown in LB medium for 2 h (from −2 h– 0 h) and treated with 0.2% (w/v) L-arabinose at the point of 0 h.

Bacteriophages of the *Microviridae* and *Leviviridae* families are small single-stranded DNA/RNA phages that infect gram-negative bacteria and express a gene encoding a protein that can lyse host bacteria. Protein E of microvirus ΦX174 and protein A_2_ of *levivirus* Qß, derived from such genes, were shown to inhibit the function of MurA and MraY, respectively, in the PG synthesis pathway (Bernhardt, Wang, Struck, & Young, n.d.; Cui et al., 2017; Reed, Langlais, Wang, & Young, 2013; Tanaka & Clemons, 2012; Timothy Bugg, Rodolis, Mihalyi, & Jamshidi, 2016) (Figure 1A). Such proteins inhibit the functions of essential enzymes involved in PG biogenesis, achieving cell lysis without directly degrading PG. Lys^M^, one such protein encoded by a lysis gene of *Levivirus* phage M (*lys*^*M*^), consists of 37 amino acid residues with a single transmembrane domain (Rumnieks & Tars, 2012). Lys^M^ prevents the lipid II flip activity of *Escherichia coli* MurJ, leading to cell lysis (Chamakura et al., 2017).

MurJ is a membrane protein consisting of 14 transmembrane helices (TMs). The TMs are divided into three parts: the N-lobe (TM1–6), C-lobe (TM7–12), and TM13–14 region. The N- and C-lobes form V-shaped structures with a central cavity for substrate binding. Structural and biological studies of MurJ implied that MurJ transports lipid II from the cytoplasm to the periplasm via its rocker switch-like conformational changes between the inward (-facing) and outward (-facing) forms (Figure 1B) (Butler, Davis, Bari, Nicholson, & Ruiz, 2013; Kohga et al., 2022; Kuk, Hao, Guan, & Lee, 2019; Kuk, Mashalidis, & Lee, 2017; Kumar, Rubino, Mendoza, & Ruiz, 2019; Rubino, Kumar, Ruiz, Walker, & Kahne, 2018; Zheng et al., 2018). In the first state, MurJ is in an inward-open form in which lipid II enters the central cavity. After lipid II binding, MurJ changes its conformation to an inward-occluded form due to lipid II headgroup interactions with the inside of MurJ. Then, the conformation changes to an outward form to release lipid II into the periplasm. After lipid II release, MurJ returns to its inward-open form for successive lipid II transport via a squeezed form. TM7, which overhangs the N-lobe from the C-lobe, is proposed to be involved in both the inward-to-outward and outward-to-inward conformational changes (Figure 1B) (Kohga et al., 2022; Kuk et al., 2019). According to the substituted cysteine accessibility method, Lys^M^ was proposed to block alternative conformational transitions of MurJ during lipid II transport. Although Lys^M^ possibly interacts with MurJ, the inhibition mechanism remains unknown.

Here, we conducted a comprehensive alanine-scanning mutagenesis study of Lys^M^ and identified crucial residues for its function. In addition, we mapped the key residues of Lys^M^ on an α-helical model, showing putative surfaces interacting with MurJ. Our findings provide basic essential information for understanding the inhibition mechanism of Lys^M^.

## Results and discussion

### Identification of amino acids crucial for Lys^M^ function

Rumnieks and Tars in 2012 determined the genome sequence of several *Leviviridae* phages. To explore the lysis gene on the *levivirus* phage M genome, they induced the expression of phage M proteins from plasmids containing cDNA copies for sequencing, and then *lys*^*M*^ was identified. Further, Chamakura et al. in 2017 cloned *lys*^*M*^ into a multicopy plasmid and analyzed the effects of Lys^M^ production on *E. coli* cells. First, we cloned *lys*^*M*^ into the pBAD33 vector. Lys^M^ expression was under the control of an arabinose-inducible promoter. Next, we prepared *E. coli* BL21 (DE3) harboring a plasmid encoding Lys^M^ to assess its toxicity in *E. coli* cells. The induction of Lys^M^ by L-arabinose decreased the density of *E. coli* cells, leading to cell death and cell lysis (Figure 1C), consistent with previous findings (Chamakura et al., 2017; Rumnieks & Tars, 2012). Because Lys^M^ is proposed to be a single inner membrane-spanning protein with an N_out_–C_in_ orientation (Chamakura et al., 2017) (Figure 1A), we introduced C-terminal tags (PA and GFP-His_6_) to Lys^M^ to detect its accumulation.

To identify crucial residues of Lys^M^, we performed comprehensive mutation analysis using alanine-substituted Lys^M^ mutants. We used 32 Lys^M^-PA mutants in which each amino acid, excluding the first methionine and four alanine residues (M1, A8, A16, A21, and A29), was replaced by alanine. The lysis activities of these mutants were tested using the spot assay on an LB plate containing L-arabinose. *E. coli* cells producing wild-type (WT) Lys^M^-PA could not grow on arabinose-supplemented LB plates or in a liquid medium similarly as cells expressing nontagged Lys^M^ (Figures 1C and 2A). Among the 32 alanine-substituted Lys^M^, 6 Lys^M^ mutants (N6A, L13A, L14A, D18A, I20A, and P30A) did not induce cell lysis, allowing cell growth even in the presence of arabinose (Figure 2A). Lys^M^ (K2A or V11A)-producing cells grew slightly in the presence of arabinose. These results identified six residues (N6, L13, L14, D18, I20, and P30) crucial for Lys^M^ function. We then predicted the Lys^M^ secondary structure and transmembrane region using the PSIPRED server and TMHMM v2.0 server, respectively. The critical residues of Lys^M^, excluding P30, were localized in the transmembrane domain (Figure 2B). Previous studies showed that Lys^M^ binds to MurJ, inhibiting the conformational changes of MurJ during the lipid II flipping cycle (Chamakura et al., 2017). Thus, the five transmembrane residues (N6, L13, L14, D18, and I20) might be involved in the interaction with MurJ or the stability of Lys^M^ itself in the membrane. Further, the helical wheel projections (Gautier, Douguet, Antonny, & Drin, 2008) of Lys^M^ showed that the essential residues of Lys^M^ create two interaction surfaces around residues 14–18 and 6–20 in the TM (Figure 2C).

**Figure 2.**
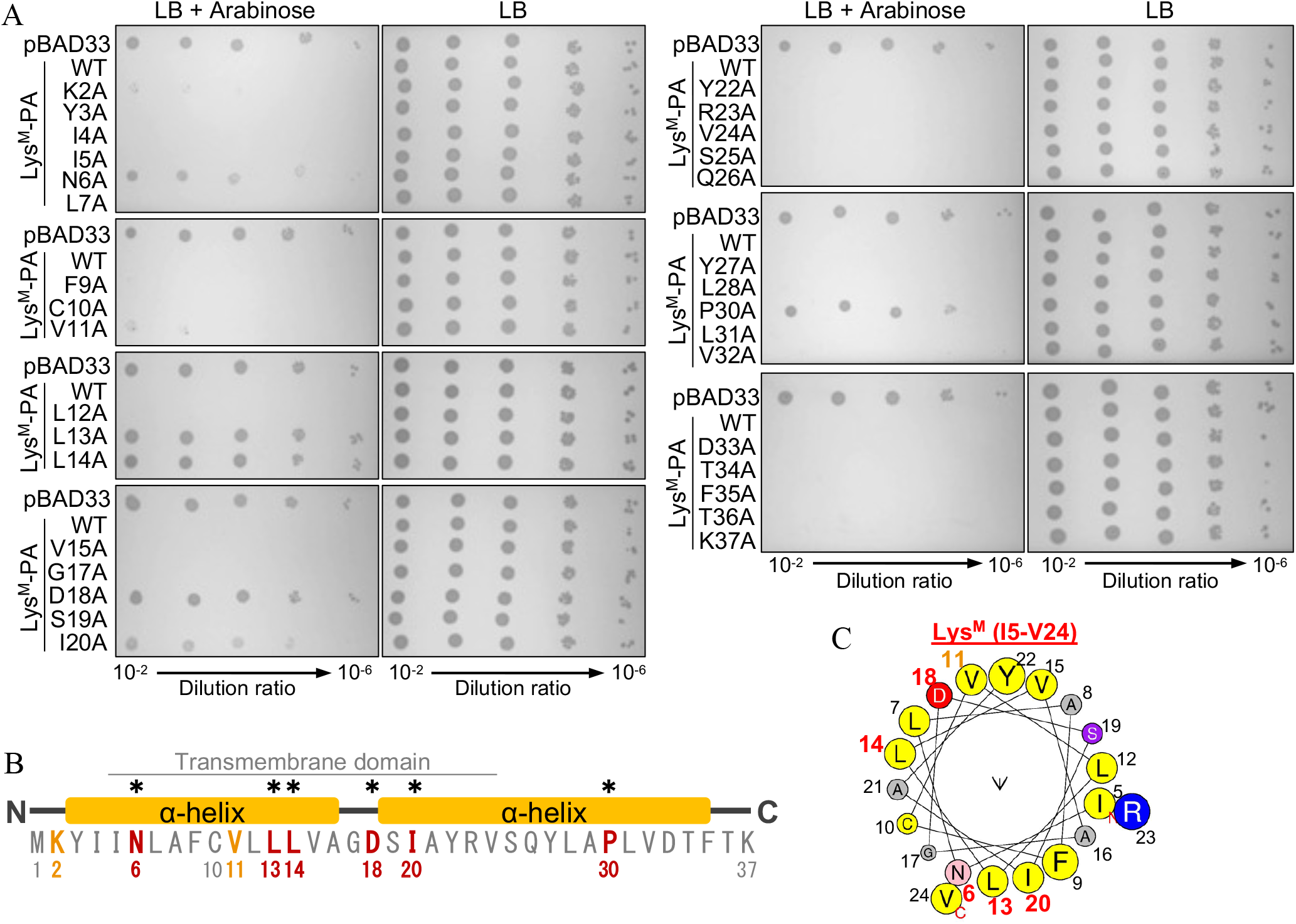
Essential residues of Lys^M^ for its cell lysis activity. (A) Effects of alanine-substituted Lys^M^ mutants on *Escherichia coli* growth. *E. coli* cells harboring pBAD33 (vector) or plasmids encoding Lys^M^ (or its mutants) were grown on LB agar plates containing 0.2% (w/v) L-arabinose. (B) Predicted secondary structure of Lys^M^ with its amino acid sequence. Asterisks indicate essential residues for the cell lysis activity of Lys^M^. The essential and important residues are highlighted in red and orange, respectively. (C) Helical wheel plot of the region predicted to form transmembrane helices of Lys^M^. The view from the periplasmic side. A helical wheel plot was generated using HELIQUEST software (https://heliquest.ipmc.cnrs.fr/, Gautier, Douguet, Antonny, & Drin, 2008). Color coding for residues was as follows: yellow for hydrophobic, purple for serine (S), blue for arginine (R), red for acidic residues, pink for asparagine (N), and gray for small residues (alanine [A] and glycine [G]). Arrows represent the magnitude and orientation of the mean hydrophobic moment value calculated by HELIQUEST software.

### The hydrophobic face on the TM is essential for the cell lysis activity of Lys^M^

We cannot exclude the possibility that the failure of cell lysis in several Lys^M^ mutants is attributable to lower protein accumulation. Therefore, we examined the accumulation of Lys^M^ mutants. Because it was challenging to handle Lys^M^-PA because of its smaller molecular mass, we also prepared a larger Lys^M^ derivative (Lys^M^-GFP-His_6_) that retained similar lysis activity as nontagged Lys^M^ (Figure 1C). To detect Lys^M^ accumulation in cells, we grew *E. coli* cells in the presence of Lys^M^. Lys^M^-induced growth deficiency in *E. coli* was reported to be complemented by the expression of BsAmj, which is a *Bacillus subtilis* lipid II flippase different from MurJ, (Adler et al., 2023; Chamakura et al., 2017; Meeske et al., 2015). Coexpression of BsAmj from a pTVW228-based plasmid blocked Lys^M^-induced cell lysis, allowing Lys^M^- producing cells to grow (Figure 3A). Using this strategy, we successfully detected Lys^M^-PA and Lys^M^- GFP-His_6_ derivatives by immunoblotting (Figure 3C and Supplementary Figure 1). In fact, it was easier to detect Lys^M^-GFP-His_6_ protein than Lys^M^-PA.

**Figure 3.**
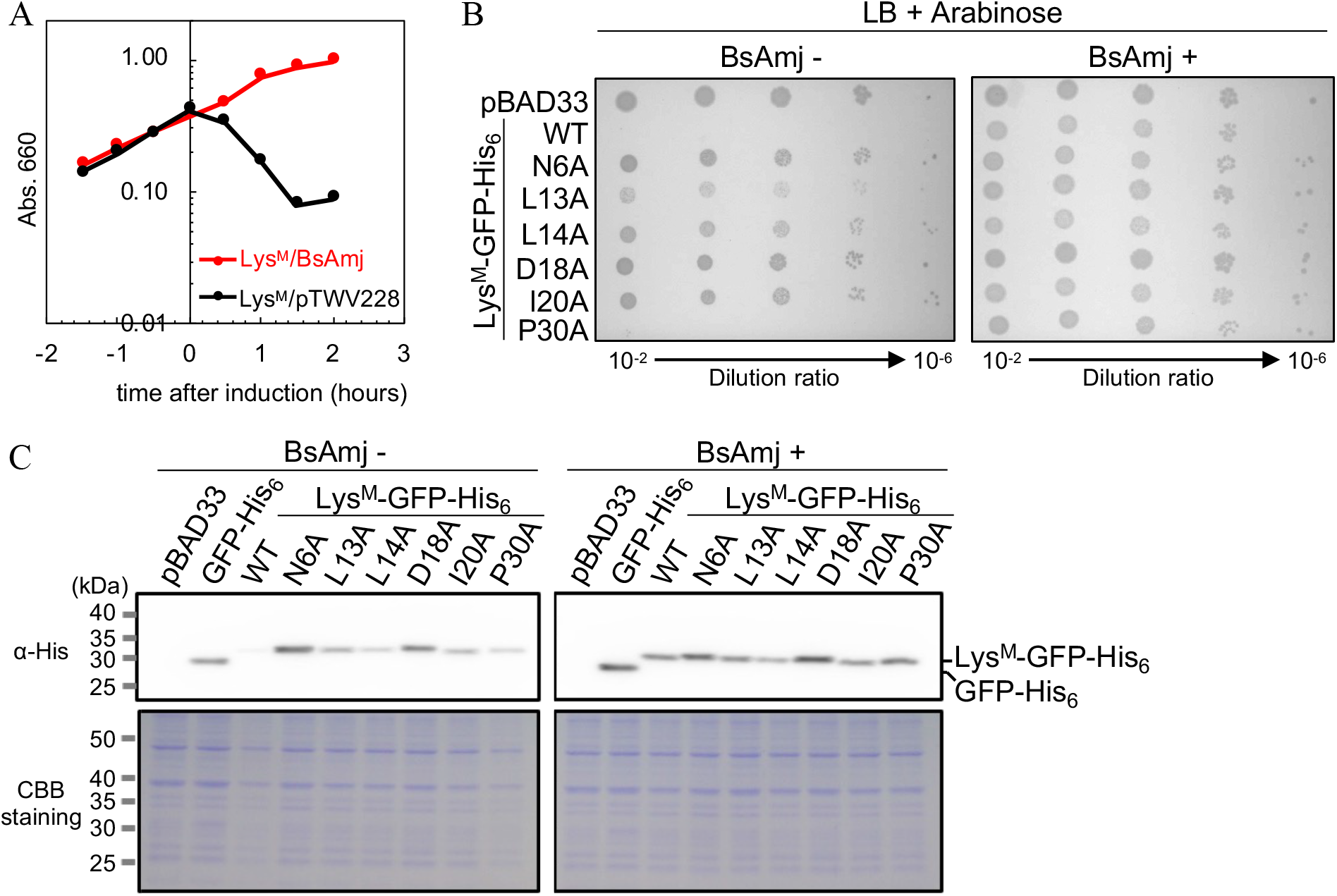
Detection of alanine-substituted Lys^M^-GFP-His_6_ mutants. (A) Suppression of Lys^M^-induced *Escherichia coli* cell lysis by BsAmj. The growth of *E. coli* cells harboring a plasmid encoding both untagged Lys^M^ and pTWV228 (vector) or a plasmid encoding BsAmj. Cells were grown in LB medium for 1.5 h and treated with 0.2% (w/v) L-arabinose and 0.2 mM IPTG at the point of 0 h. (B) Spot assay of *E. coli* cells expressing both alanine-substituted Lys^M^-GFP-His_6_ and BsAmj. Cells were grown on LB agar plates containing 0.2% (w/v) L-arabinose. (C) Accumulation of Lys^M^- GFP-His_6_ in each transformant. Cells were grown in LB medium until the early log-phase and treated with 0.2% (w/v) L-arabinose and 0.2 mM IPTG for 2 h. After induction, total cell lysate samples were acid-precipitated and analyzed using SDS-PAGE and immunoblotting with anti-His antibody.

Comparing the cell lysis activities of mutant and WT Lys^M^-GFP-His_6_ on an arabinose-supplemented LB plate, Lys^M^-GFP-His_6_ mutants, excluding P30A, did not induce cell lysis similarly as Lys^M^-PA in the presence of arabinose (Figure 3B left). By contrast, the additional expression of Amj enhanced the growth of all cells expressing Lys^M^-GFP-His_6_ (Figure 3B right). All Lys^M^-GFP-His_6_ constructs, including WT, were detected upon Amj expression, and their accumulation levels were comparable (Figure 3C right). Even in the absence of Amj expression, Lys^M^ (N6A, L13A, L14A, D18A, I20A)-GFP-His_6_ was confirmed to be sufficiently accumulated (Figure 3C left).

Although the expression of WT Lys^M^-GFP-His_6_ induced cell lysis in the absence of Amj, we detected a small amount of WT Lys^M^-GFP-His_6_ in the medium after cell lysis. Conversely, it was unclear why Lys^M^ (P30A)-GFP-His_6_ had cell lysis activity (Figure 3B), contradicting the results for Lys^M^ (P30A)- PA (Figure 2A). Other Lys^M^ (N6A, L13A, L14A, D18A, and I20A)-GFP-His_6_ mutants were stably accumulated without inducing cell lysis, implying that these mutations affect their interaction with MurJ. Based on the helical wheel plot, the five residues of Lys^M^ were located on two surfaces of the α-helix model (Figure 2C). The two surfaces might provide the interaction site for MurJ. Because leucine and isoleucine differ from alanine only in the length of the carbon chain, an adequate hydrophobic interaction between LysM and MurJ in the membrane could be indispensable.

We then considered the inhibition mechanism of Lys^M^, which locks the conformational change of MurJ (Chamakura et al., 2017), based on structural studies of MurJ. The TM7 region of MurJ plays an important role in both inward-to-outward and outward-to-inward conformational changes (Kohga et al., 2022; Kuk et al., 2019) (Figure 1B). TM7 features a straight architecture in the outward form of TaMurJ (PDB ID: 6NC9), whereas it has a bending architecture in the squeezed form, which is an intermediate form during the transition from outward-to-inward forms of EcMurJ (PDB ID: 7WAG). Dynamic TM7 rearrangements were found to be important for both conformational changes and function. In addition, screening for Lys^M^-resistant mutants revealed that single-residue substitutions in MurJ were mapped in TM2 and TM7 (Chamakura et al., 2017). In particular, the resistant mutations in TM7 were located on the surface in both the inward and outward structures of MurJ, and these residues might interact with Lys^M^. Lys^M^ could bind near TM7 in MurJ, inhibiting conformational changes in TM7. Recently, it was reported that a small lysis protein (Sgl^PP7^) in the ssRNA phage PP7, which infects *Pseudomonas aeruginosa*, also targets MurJ (Adler et al., 2023). Furthermore, a suppression mutation of Sgl^PP7^ is also located in TM7 of MurJ (Adler et al., 2023). The mechanism of inhibition of Sgl^PP7^ might be similar to that of Lys^M^.

In this study, we revealed that key residues of Lys^M^ cluster in a TM, suggesting that these residues may form the interaction site for MurJ. Therefore, our findings provide clues for understanding the working mechanism of Lys^M^. However, the direct interaction between Lys^M^ and MurJ and the actual interaction sites remain unclear. To further understand the mechanism of Lys^M^ inhibition, structural analysis of the MurJ/Lys^M^ complex is required. In addition, studies of small lysis proteins of ssRNA phages will provide helpful information for developing antimicrobial peptides.

## Author contributions

Conceptualization, H.K., T.T.; Methodology and Investigation, H.K., N.L., R.M., T.T.; Writing – Original Draft, H.K., T.T.; Writing – Review and Editing, H.K., T.T.; Supervision, T.T.

## Materials and Methods

### Plasmids and E. coli cells

Plasmids used in this study are listed in Supplemental Table 1.

A DNA fragment containing *lys*^*M*^ gene and its Shine-Dalgarno sequence, 5’- CTCGGTACCCGGGGATCCTCTAGGAGGTTTAAATTATGAAATATATAATAAATTTAGCATTCTGCGTTCTCTTACTGGTTGCGGGGGACTCGATAGCATATCGAGTCTCGCAATACCTGGCGCCTTTGGTGGATACCTTCACCAAGTAAAGAGTCGACCTGCAGGCATGC-3’, and XbaI-digested pBAD33 were changed into a plasmid by Gibson assembly (Gibson et al., 2009). The resulting plasmid was named pKK568. A DNA, 5’-GGAGTCGCCATGCCCGGAGCCGAGGATGATGTCGTC-3’, encoding PA tag was inserted before the stop codon of *lys*^*M*^ and the resulting plasmid was named pKG74. The plasmids with mutations as shown in Supplemental Table1 were used. Modified plasmids (pKG74– pKG106) based on pKG74 are shown in Supplemental Table 1.

For the construction of plasmids for Lys^M^-GFP-His_6_ expression, the *gfp-his6* gene was PCR amplified from pCGFP-BC (Kawate & Gouaux, 2006) using primers GFP-His6_f (5’- CTTTGGTGGATACCTTCACCAAGGGCGGTAGCGGCCAATTTACTAGTAGCGTGAGTAAAG-3’) and GFP-His6_r (5’- AACAGCCAAGCTTGCATGCCTCAGTGATGGTGATGGTGATGGTGA-3’). The PCR product was cloned into SalI-PstI-digested pKG120 by In-Fusion HD Cloning Kit (Takara bio). The resulting plasmid, pKG126, was used as templates of site-directed mutagenesis for making pNL1, 4, 7, 10, 13, and 15.

For the construction of pRM1071, the *Bacillus subtilis amj* fragment was PCR amplified from the genome of *Bacillus subtilis subsp. subtilis* JCM 1465T (RIKEN BRC) using a pair of primers, Bs_amj-F (5’-GCGCGAATTCATTGGAGGAAGAATAACGTGC-3’) and Bs_amj-R (5’- GCGCAAGCTTTTAAAACCACTTTGTCAGCC-3’). The PCR product was digested with EcoRI and HindIII, and ligated to the corresponding sites in pTWV228 (Takara bio). The resulting plasmid was named pRM1071.

For the construction of pRM1210, PCR amplified DNA fragment containing the *Bacillus subtilis amj* using a pair of primer, Bs_amj-F2 (5’-CAGGAAACAGCCATGCATGTCATTACAACACAAG-3’) and Bs_amj-R2 (5’-GGCCAGTGCCAAGCTTTTAAAAC-3’), from pRM1071 was cloned into NcoI-HindIII-digested pHM1552 (Miyazaki et al., 2022) using In-Fusion HD Cloning Kit (Takara bio).

### Growth conditions and Strains

*E. coli* cells were grown at 37 °C in LB media, supplemented with chloramphenicol (50 μg/mL) or ampicillin (100 μg/mL). For the cell lysis activity test, *E. coli* BL21 (DE3) strains (Laboratory stock) were transformed with pBAD33 or pKG74–106. Transformants were grown in LB media for 2 hours and induced with 0.2% (w/v) L-arabinose. Growth was monitored by optical density at 660 nm. For Immunoblotting analysis, BL21 (DE3) cells were transformed with a pair of plasmids pRM1072 encoding *Bacillus subtilis* Amj and encoding Lys^M^–GFP-His_6_ or alanine-substituted Lys^M^-GFP-His_6_ (pKG164 or plasmids pNL1–15).

### Spot-dilution plate assay

Cells in LB media, supplemented with appropriate antibiotics, were grown overnight were diluted 1: 100 in PBS. The diluted cultures have serially diluted ten-fold, and 5 μL of dilutions were spotted onto LB agar plates supplemented with 0.2% (w/v) L-arabinose and appropriate antibiotics. The plates were incubated at 37 °C and imaged after 22 hours.

### Immunoblotting analysis

Lys^M^-GFP-His_6_- and BsAmj-expressing cells were grown in LB media with appropriate antibiotics, until early log-phase, and induced using 0.2% L-arabinose and 0.2 mM IPTG for 2 hours. Total cellular proteins precipitated by TCA were solubilized in a SDS sample buffer (62.5 mM Tris-HCl (pH 8.0, 2% SDS, 10% glycerol, 5% β-mercaptoethanol), and separated by SDS-PAGE using 10% Laemmli gel. For detection of the Lys^M^-PA protein, total cellular proteins were solubilized in the SDS sample buffer, and analyzed using 15% Bis-Tris gel and MES SDS running buffer as described previously (Yokoyama et al., 2021). Proteins were transferred onto PVDF membranes (Merck Millipore). The membranes were blocked with skim milk, and probed with rabbit anti-His primary antibody (1: 5,000 dilution, MBL PM032) and, goat anti-rabbit IgG HRP-conjugated secondary antibody (1: 3,000 dilution, BIO-RAD 170-6515). Proteins were detected with Chemi-Lumi One (Nacalai Tesque) using FUSION Solo S (VILBER).

**Supplemental Figure 1 Accumulation check of Lys^M^-PA**

Accumulation of Lys^M^-PA in *Escherichia coli* cells harboring pBAD33 (vector) or plasmids encoding Lys^M^-PA or BsAmj (pRM1210). Lys^M^-PA was detected as described in Figure 3C, except for the use of anti-PA antibody.

## Acknowledgments

We thank Kayo Abe for secretarial assistance; Kumi Kobayashi for construction of pKK568; and Enago (www.enago.jp) for the English language review. This work was supported by JSPS/MEXT KAKENHI (Grant No. JP23K14146 to K.G., Grant No. JP22K15061, JP22H05567 to R.M., and Grant Nos. JP22H02567, JP22H02586, JP21H05155, JP21H05153, JP21K19226, JP21KK0125 to T.T.), and private research foundations (the Chemo-Sero-Therapeutic Research Institute, Naito Foundation, Takeda Science Foundation, G-7 Scholarship Foundation, the Sumitomo Foundation, the Institute for Fermentation (Osaka), Yamada Science Foundation, and Japan Foundation for Applied Enzymology) to T.T. The genome of *Bacillus subtilis subsp. subtilis* JCM 1465T was provided by RIKEN BRC, which is participating in the National Bio-Resources Project of MEXT, Japan.

