## Supplemental Table 1 and Supplemental Figure 1 for "Critical residues of the antibiotic peptide Lys^M^ that inhibits lipid II flipping"

| Plasmids | Vector | Genotype | Reference |
|---|---|---|---|
| pBAD33 | - | Expression vector, $P_{araBAD}$, $Cm^R$ | Takara Bio |
| pKK568 | pBAD33 | *lys*$^M$ | This study |
| pKG74 | pBAD33 | *lys*$^M$*-pa* | This study |
| pKG75 | pBAD33 | *lys*$^M$*(K2A)-pa* (by mutating AAA to GCA) | This study |
| pKG76 | pBAD33 | *lys*$^M$*(Y3A)-pa* (by mutating TAT to GCA) | This study |
| pKG77 | pBAD33 | *lys*$^M$*(I4A)-pa* (by mutating ATA to GCA) | This study |
| pKG78 | pBAD33 | *lys*$^M$*(I5A)-pa* (by mutating ATA to GCA) | This study |
| pKG79 | pBAD33 | *lys*$^M$*(N6A)-pa* (by mutating AAT to GCA) | This study |
| pKG80 | pBAD33 | *lys*$^M$*(L7A)-pa* (by mutating TTA to GCA) | This study |
| pKG81 | pBAD33 | *lys*$^M$*(F9A)-pa* (by mutating TTC to GCA) | This study |
| pKG82 | pBAD33 | *lys*$^M$*(C10A)-pa* (by mutating TGC to GCA) | This study |
| pKG83 | pBAD33 | *lys*$^M$*(V11A)-pa* (by mutating GTT to GCA) | This study |
| pKG84 | pBAD33 | *lys*$^M$*(L12A)-pa* (by mutating CTC to GCA) | This study |
| pKG85 | pBAD33 | *lys*$^M$*(L13A)-pa* (by mutating TTA to GCA) | This study |
| pKG86 | pBAD33 | *lys*$^M$*(L14A)-pa* (by mutating CTG to GCA) | This study |
| pKG87 | pBAD33 | *lys*$^M$*(V15A)-pa* (by mutating GTT to GCA) | This study |
| pKG88 | pBAD33 | *lys*$^M$*(G17A)-pa* (by mutating GGG to GCA) | This study |
| pKG89 | pBAD33 | *lys*$^M$*(D18A)-pa* (by mutating GAC to GCA) | This study |
| pKG90 | pBAD33 | *lys*$^M$*(S19A)-pa* (by mutating TCG to GCA) | This study |
| pKG91 | pBAD33 | *lys*$^M$*(I20A)-pa* (by mutating ATA to GCA) | This study |
| pKG92 | pBAD33 | *lys*$^M$*(Y22A)-pa* (by mutating TAT to GCA) | This study |
| pKG93 | pBAD33 | *lys*$^M$*(R23A)-pa* (by mutating CGA to GCA) | This study |
| pKG94 | pBAD33 | *lys*$^M$*(V24A)-pa* (by mutating GTC to GCA) | This study |
| pKG95 | pBAD33 | *lys*$^M$*(S25A)-pa* (by mutating TCG to GCA) | This study |
| pKG96 | pBAD33 | *lys*$^M$*(Q26A)-pa* (by mutating CAA to GCA) | This study |
| pKG97 | pBAD33 | *lys*$^M$*(Y27A)-pa* (by mutating TAC to GCA) | This study |
| pKG98 | pBAD33 | *lys*$^M$*(L28A)-pa* (by mutating CTG to GCA) | This study |
| pKG99 | pBAD33 | *lys*$^M$*(P30A)-pa* (by mutating CCT to GCA) | This study |
| pKG100 | pBAD33 | *lys*$^M$*(L31A)-pa* (by mutating TTG to GCA) | This study |
| pKG101 | pBAD33 | *lys*$^M$*(V32A)-pa* (by mutating GTG to GCA) | This study |
| pKG102 | pBAD33 | *lys*$^M$*(D33A)-pa* (by mutating GAT to GCA) | This study |
| pKG103 | pBAD33 | *lys*$^M$*(T34A)-pa* (by mutating ACC to GCA) | This study |
| pKG104 | pBAD33 | *lys*$^M$*(F35A)-pa* (by mutating TTC to GCA) | This study |
| pKG105 | pBAD33 | *lys*$^M$*(T36A)-pa* (by mutating ACC to GCA) | This study |
| pKG106 | pBAD33 | *lys*$^M$*(K37A)-pa* (by mutating AAG to GCA) | This study |
| pKG126 | pBAD33 | *lys*$^M$*-gfp-*$his_6$ | This study |
| pKG164 | pBAD33 | *gfp-*$his_6$ | This study |
| pNL1 | pBAD33 | *lys*$^M$*(N6A)-gfp-*$his_6$ (by mutating AAT to GCA) | This study |
| pNL4 | pBAD33 | *lys*$^M$*(L13A)-gfp-*$his_6$ (by mutating TTA to GCA) | This study |
| pNL7 | pBAD33 | *lys*$^M$*(L14A)-gfp-*$his_6$ (by mutating CTG to GCA) | This study |
| pNL10 | pBAD33 | *lys*$^M$*(D18A)-gfp-*$his_6$ (by mutating GAC to GCA) | This study |
| pNL13 | pBAD33 | *lys*$^M$*(I20A)-gfp-*$his_6$ (by mutating ATA to GCA) | This study |
| pNL15 | pBAD33 | *lys*$^M$*(P30A)-gfp-*$his_6$ (by mutating CCT to GCA) | This study |
| pTWV228 | - | Expression vector, $P_{lac}$, $Amp^R$ | Takara Bio |
| pRM1072 | pTWV228 | *Bacillus subtilis amj* | This study |
| pRM1210 | pMW118 | *Bacillus subtilis amj* | This study |

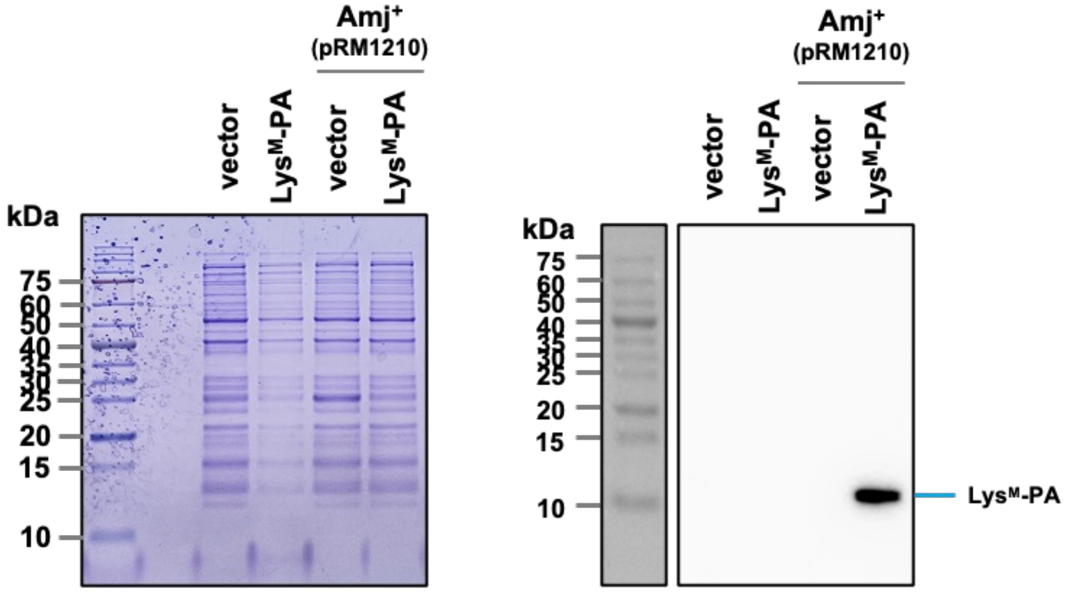


**Supplemental Figure 1 Accumulation check of Lys$^{M}$-PA**
Accumulation of Lys$^{M}$-PA in *Escherichia coli* cells harboring pBAD33 (vector) or plasmids encoding Lys$^{M}$-PA or BsAmj (pRM1210). Lys$^{M}$-PA was detected as described in Figure 3C, except for the use of anti-PA antibody.
